# The influence of captivity on cardiac structure and function across age, in rhesus macaques

**DOI:** 10.1101/2025.10.20.683517

**Authors:** T G Dawkins, B A Curry, A Drane, V N Rivas, Y Ueda, J A Stern, D Philips, J Negron-Del Valle, Cayo Biobank Research Unit, JP Higham, N Snyder-Mackler, L Brent, R Shave

**Affiliations:** Centre for Heart, Lung, and Vascular Health, School of Health and Exercise Sciences, University of British Columbia Okanagan, Kelowna, Canada; Faculty of Medicine, Health & Life Sciences, Swansea University, Swansea, United Kingdom; Department of Clinical Sciences, College of Veterinary Medicine, North Carolina State University, Raleigh, North Carolina, United States; California National Primate Research Center, University of California, Davis, Davis, California, United States; School of Life Sciences, Arizona State University, Tempe, Arizona, United States; Department of Anthropology, New York University, New York, New York, United States; Centre of Research in Animal Behaviour, University of Exeter, Exeter, United Kingdom

## Abstract

**Background:** Captive non-human primates are widely used as models of human aging, yet the conditions they live in differ markedly from their naturalistic environment. Differences between captive and free-ranging environments may impact how the cardiovascular system adapts with age, potentially confounding studies of natural aging. This study characterized age-related cardiac phenotypes in free-ranging rhesus macaques and compared these patterns with their captive-housed counterparts to assess the influence of living environment on cardiac health across the lifespan.

**Methodology:** We performed transthoracic echocardiography in a cross-sectional cohort of 133 free-ranging rhesus macaques (*Macaca mulatta*, aged 7 - 25 years, 41 % female) living on Cayo Santiago, Puerto Rico. First, we compared structural and functional cardiac parameters between young (7 - 12 years; n = 48, 60% female) and old (18-26 years, n = 33, 42% female) free-ranging macaques. We then performed an ANCOVA, with age as a covariate, to compare cardiac structural and functional parameters between free-ranging and captive-housed macaques (n = 347, aged 7 - 32 years, 71% female).

**Results:** In our free-ranging cohort, older macaques had greater interventricular septal thickness and relative wall thickness (RWT) than young. Males exhibited larger left ventricular (LV) internal dimensions, wall thickness, LV mass, and LV volumes than females, but these differences were attenuated when indexed to body mass. Diastolic function was lower with advanced age in both sexes, reflected by a lower E/A ratio, reduced myocardial tissue velocities during early diastole (e′) and greater myocardial tissue velocities during atrial contraction (a’). Compared with captive macaques, free-ranging animals exhibited better diastolic function, including a significantly higher E/A ratio, greater e’, and lower a’. Captive macaques also had significantly greater relative wall thickness across age and between sexes.

**Conclusions and implications:** Our study provides the first comprehensive characterization of age-related cardiac differences in free-ranging rhesus macaques, which show structural and functional cardiac differences similar to those observed with human aging. Captive macaques exhibited a more pronounced age-related cardiac phenotype than their free-ranging counterparts, including thicker left ventricular walls and lower diastolic function for a given age. These findings highlight the importance of considering the ecological context when interpreting animal models of cardiovascular aging.

## Introduction

Human aging is characterized by a gradual decline in physiological function (1). Central to this systemic decline is a progressive deterioration of the cardiovascular (CV) system (2,3), which includes an increase in left ventricular (LV) wall thickness and reduced diastolic function (1) - hallmark indices of cardiac aging. Indeed, findings from both the Framingham Heart Study, and the UK Biobank indicate that with each progressive decade of life, the ratio of early to late diastolic ventricular filling velocity (E/A ratio) decreases (4), alongside a concurrent decline in diastolic myocardial deformation (5) and progressive structural remodeling, including greater ventricular wall thickness (6). Although such studies are highly informative, the data are often confounded by numerous lifestyle factors, such as diet, physical activity, smoking, access to health care, and social stressors (e.g., social adversity, financial stress). Such confounding factors contribute to significant heterogeneity within these cohorts, complicating efforts to disentangle the direct effects of chronological aging from those of lifestyle and environmental influences on the cardiac phenotype.

In an attempt to better understand the aging cardiac phenotype in a highly controlled setting, many researchers have turned to animal models, such as rodents and non-human primates (7,8). For example, Florio and colleagues recently observed age-related differences in the cardiac phenotype between young and old captive rhesus macaques similar to that seen in humans, inferring a decline in diastolic function (9). Although animal models can minimize the impact of some confounding variables, it is important to acknowledge that these animals live in a highly artificial environment, which is markedly different from their evolutionary niche. Findings from captive animal populations may also be confounded by lifestyle factors. Previous work has shown that, compared with free-ranging animals, captive non-human primates have a greater body mass (10), proatherogenic lipid profiles (11), physiological dysregulation (12), and a higher prevalence of cardiovascular disease (13,14). Accordingly, differences between young and old animals may not solely reflect an age-related decline, but also the potential interaction between the artificial environment and advancing age.

To further our understanding of cardiac aging in free-living macaques, and to explore the influence of the captive environment, we undertook a study with two aims. First, we characterized cardiac structure and function in young and old, male and female, adult free-ranging rhesus macaques. Second, we compared the cardiac phenotype between free-ranging and captive rhesus macaques across the adult lifespan. We predicted that: i) in free-ranging rhesus macaques, advancing age would be associated with thicker ventricular walls and lower diastolic function; and ii) captivity would be associated with a more pronounced age-related cardiac phenotype.

## Methods

### Rhesus Macaque Cohorts

Echocardiographic assessments were performed in free-ranging (n = 133) and captive (n = 422) adult rhesus macaques >7 years of age. Free-ranging macaques inhabited the island of Cayo Santiago, Puerto Rico, managed by the Caribbean Primate Research Center. Cayo Santiago is home to ∼1700 free-ranging rhesus macaques, which are descendants of 409 rhesus macaques transported from India in 1938. Animals are provisioned with monkey chow and have *ad libitum* access to drinking water. The monkeys live in naturally forming social groups characterized by dominance hierarchies and male dispersal. The island is free of predators and there is no regular veterinary intervention. Free-ranging animals were lured into a corral and captured with a net, and were subsequently transferred to a squeeze cage, and were anesthetized with intramuscular ketamine HCL (100mg/mL solution at 1.0 mL and 0.8 mL for males and females, respectively) and intravenous xylazine (100 mg/ml solution at 0.05mL and 0.03mL for males and females, respectively). Supplemental ketamine and xylazine were given as an intravenous bolus when required to maintain anaesthesia.

The data from captive animals were obtained from a previously published database, taken at the US Davis California National Primate Research Centre (15,16) and managed in accordance with the Animal Welfare Act and the Guide for the Care and Use of Laboratory Animals. Outdoor-housed animals lived in social groups within 0.5-acre enclosures, while indoor-housed animals were pair- or singly housed in stainless steel cages meeting space requirements. All animals received environmental enrichment, a standard primate chow diet supplemented with fruits and vegetables, and *ad libitum* access to water. Veterinary staff conducted routine health monitoring, including physical examinations, hematology, biochemistry, tuberculosis testing, and viral screening. Animals housed in captivity were anesthetized using ketamine, as described previously (15,16).

Research in free-ranging animals was approved by the University of British Columbia Animal Care Committee (ID: A22-0180) and Institutional Animal Care and Use Committee (IACUC; ID: A400117). Protocols involving captive animals were approved by IACUC (base grant #P51OD011107) at the UC Davis California National Primate Research Centre (15,16). In line with data available from our free-ranging population, only captive animals >7 years of age were included. No animals were euthanized for the completion of this study.

### Transthoracic Echocardiographic Assessments

Echocardiography assessments in the free-ranging population were completed using a commercially available ultrasound machine (Vivid IQ, GE Healthcare, Chalfont St Giles, Bucks, UK) with a 5.0 – 11.0 MHz phased-array transducer. All cardiac examinations in free-ranging macaques were conducted by an experienced research sonographer (TD). Three cardiac cycles were collected for each animal, and stored for offline analysis (EchoPac, GE Medical, Horten, Norway, version 204). Indirect brachial blood pressures were obtained using an automated system (SphygmoCor Xcel, AtCor Medical, Sydney, Australia) at the onset of the echocardiographic assessment. Measurements of cardiac structure included aortic annulus dimension, septal wall thickness (IVSDd), posterior wall thickness (PWDd), internal LV dimensions (LVIDd) and relative wall thickness (RWT; the sum of septal and posterior wall thickness, divided by LV internal dimension), obtained from the parasternal long axis. LV mass was calculated using the Cube formula at the level of the parasternal long axis (17). LV sphericity index was calculated from the apical four-chamber view as LV length/diameter during diastole. Biplane LV volumes were analyzed using the automated functional imaging software, tracing the endocardial border at end-diastole and end-systole from the apical four-and apical two-chamber views. SV was calculated by subtracting end-systolic volume from end-diastolic volume (EDV). Given the known relationship between body mass and cardiac structure, parameters were scaled, where appropriate, to body mass raised to an allometric exponent based on the model of quarter-power scaling (18). LV diastolic function was assessed by measuring mitral inflow velocities during early left ventricular filling (mitral E wave) and late ventricular filling (mitral A wave) from the apical four-chamber view. Tissue Doppler velocities were also taken from the apical window at the level of the mitral annulus, for the septum and lateral LV wall, to give systolic (s’) early diastolic (e’) and late diastolic (a’) tissue velocities. Myocardial strain and strain rate (i.e., deformation of the myocardium) were assessed by tracking the LV wall throughout systole and diastole using two-dimensional speckle tracking analysis from an apical four-chamber view. Left atrial (LA) maximal volume and LA deformation were generated using the automated functional imaging speckle tracking tool in EchoPac.

Details of the echocardiographic assessments in captive macaques have been published previously (15). Briefly, echocardiography was performed using a commercial ultrasound system (Affinity 50, Phillips Amsterdam, Netherlands), with images analyzed offline using Syngo Dynamics software (Siemens, Erlangen, Germany). Animals with overt structural disease were excluded from the analysis (n = 105; primarily left ventricular hypertrophy), resulting in a cohort of 317 captive macaques. For this comparison, we specifically focused on hallmark features of cardiac aging, including relative wall thickness (RWT), the ratio of early to late left ventricular filling (E/A), and early and late diastolic tissue velocities (e’ and a’).

### Statistical analysis

A two-way ANOVA was conducted to assess the main effects of age (young vs. old) and sex (male vs. female), as well as their potential interaction, in the free-ranging macaque population. Normality of the model residuals was assessed using the Shapiro-Wilk test and inspection of Q-Q plots. Pairwise comparisons were performed for each significant main effect, with Bonferroni-adjusted *P* values within each simple main effect. A two-way ANCOVA was then performed to examine our second aim, assessing the influence of environment (captive vs. free-ranging macaques) and sex (male vs. female) on the cardiac phenotype, incorporating age as a covariate (SPSS Statistics, Windows version 29, IBM Corp., Armonk, N.Y. USA). For E, A, and E/A ratio, heart rate was also included as a covariate. Where there was a significant age*environment*sex interaction, slopes of the regression lines were compared to assess whether the relationships between age and the dependent variables differed between captive and free-ranging macaques. All ANCOVA statistical analyses were performed using the Statistical Package for the Social Sciences (SPSS; version 29, IBM Corp., Armonk, N.Y. USA). Comparison of regression slopes were performed using GraphPad Prism (version 10.3.0, Boston, Massachusetts, USA). Data are reported as mean ± standard deviation, and statistical significance was set at α = 0.05.

## Results

Demographic characteristics and blood pressures of free-ranging macaques are reported in **Table 1**. All LV and LA parameters are reported in Table 2. Exact P values for pairwise comparisons are provided in Supplementary Table 1.

**Table 1.** Descriptive characteristics.

| Parameter | All | Female |  | Male |  |
| --- | --- | --- | --- | --- | --- |
|  |  | Young<br>(7-12 yrs) | Old<br>(18-26 yrs) | Young<br>(7-12 yrs) | Old<br>(18-26 yrs) |
| n = | 81 | 29 | 14 | 19 | 19 |
| Age (Years) | 14.5 ± 4.7 | 10.4 ± 1.5 | 19.6 ± 1.9* | 10.9 ± 1.3 | 20.0 ± 1.4* |
| Body mass (kg) | 8.6 ± 1.9 | 7.6 ± 0.7 | 6.6 ± 0.7* | 10.2 ± 1.9† | 9.2 ± 0.7*† |
| Body length (cm) | 442 ± 49 | 414 ± 20 | 425 ± 53 | 459 ± 44† | 471 ± 51† |
| SBP brachial (mmHg) | 94 ± 17 | 100 ± 19 | 90 ± 15* | 96 ± 17 | 88 ± 13 |
| DBP brachial (mmHg) | 57 ± 14 | 59 ± 17 | 52 ± 10 | 62 ± 14 | 53 ± 10* |
| Heart rate (bpm) | 86 ± 17 | 90 ± 26 | 84 ± 14 | 84 ± 11 | 86 ± 16 |

\* *P* < 0.05 vs young, within sex; † *P* < 0.05 vs female, within age group.

**Table 2.** Cardiac structure and function in male and female, young and old free-ranging rhesus macaques.

| Echo Parameter | All | Female |  | Male |  |
| --- | --- | --- | --- | --- | --- |
|  |  | Young | Old | Young | Old |
|  |  | (7-12 yrs) | (18-26 yrs) | (7-12 yrs) | (18-26yrs) |
| LV structure |  |  |  |  |  |
| LV mass (g) | 15 ± 4 | 12 ± 3 | 12 ± 4 | 17 ± 4† | 18 ± 4† |
| LV mass/body mass (g/kg<sup>1.0</sup>) | 1.7 ± 0.4 | 1.7 ± 0.3 | 1.9 ± 0.5 | 1.6 ± 0.3 | 2.0 ± 0.4* |
| Aortic annulus diameter (mm) | 10.3 ± 0.95 | 9.4 ± 0.5 | 9.7 ± 0.6 | 10.7 ± 0.6† | 11.3 ± 0.77*† |
| Aortic annulus diameter/body mass (mm/kg<sup>0.25</sup>) | 6.0 ± 0.4 | 5.7 ± 0.3 | 6.1 ± 0.3* | 6.0 ± 0.4† | 6.5 ± 0.5*† |
| IVSDd (mm) | 3.8 ± 0.8 | 3.5 ± 0.5 | 4.0 ± 0.8* | 3.7 ± 0.7 | 4.4 ± 1.2* |
| IVSDd/body mass (mm/kg<sup>0.25</sup>) | 2.2 ± 0.5 | 2.1 ± 0.3 | 2.5 ± 0.5* | 2.0 ± 0.4 | 2.5 ± 0.7* |
| LVIDd (mm) | 23.3 ± 2.7 | 22.4 ± 1.4 | 20.8 ± 2.8 | 24.7 ± 1.8† | 24.6 ± 2.5† |
| LVIDd/body mass (mm/kg<sup>0.25</sup>) | 13.6 ± 1.2 | 13.6 ± 0.7 | 12.9 ± 1.7 | 13.8 ± 0.9 | 14.1 ± 1.3† |
| PWDd (mm) | 4.1 ± 0.7 | 3.7 ± 0.6 | 3.9 ± 0.7 | 4.1 ± 0.7† | 4.1 ± 0.7† |
| PWDd/body mass (mm/kg<sup>0.25</sup>) | 2.4 ± 0.4 | 2.3 ± 0.4 | 2.4 ± 0.4 | 2.4 ± 0.3 | 2.5 ± 0.3 |
| Relative wall thickness | 0.34 ± 0.07 | 0.32 ± 0.05 | 0.39 ± 0.08* | 0.32 ± 0.05 | 0.36 ± 0.08 |
| Sphericity index | 1.61 ± 0.14 | 1.64 ± 0.14 | 1.68 ± 0.13 | 1.55 ± 0.11† | 1.59 ± 0.16† |
| LV Volumes |  |  |  |  |  |
| LV EDV biplane (ml) | 16 ± 4 | 15 ± 2 | 13 ± 3* | 19 ± 3† | 16 ± 3*† |
| LV EDV/body mass (ml/kg<sup>1.0</sup>) | 1.83 ± 0.42 | 2.01 ± 0.30 | 1.82 ± 0.57 | 1.81 ± 0.26 | 1.68 ± 0.52 |
| LV ESV biplane (ml) | 8 ± 2 | 7 ± 1 | 6 ± 1 | 9 ± 2† | 8 ± 1*† |
| LV ESV/body mass (ml/kg<sup>1.0</sup>) | 0.86 ± 0.21 | 0.94 ± 0.18 | 0.83 ± 0.27 | 0.89 ± 0.14 | 0.77 ± 0.24 |
| LV SV biplane (ml) | 8 ± 2 | 8 ± 2 | 7 ± 2 | 10 ± 2† | 9 ± 2† |
| LV SV/body mass (ml/kg<sup>1.0</sup>) | 0.96 ± 0.24 | 1.04 ± 0.16 | 0.99 ± 0.31 | 0.92 ± 0.19 | 0.92 ± 0.29 |

Abbreviations: IVSDd, intraventricular septum diameter during diastole; LVIDd, left ventricular internal diameter during diastole; PWDd, posterior wall diameter during diastole; LV, left ventricle; EDV, end diastolic volume; ESV, end systolic volume; SV, stroke volume.
\* P < 0.05 vs young, within sex; † P < 0.05 vs female, within age group.

### Characterization of cardiac phenotype in young and old free-ranging macaques

*Left ventricular structure and volume in free-ranging macaques;* males had a greater LV mass than females, which did not differ by age (**Table 2 and Supplementary Table 1**). However, when body mass was considered (i.e., LV mass scaled to body mass), no sex difference was observed in LV mass. Instead, older males had a greater scaled LV mass than their younger counterparts. Older males also had a greater aortic annulus diameter compared with their younger counterparts. In females, no age-related differences were detected until aortic diameter was scaled to body mass. In both younger and older macaques, males had a greater aortic diameter compared with females, even when scaled to body mass. IVSd and relative wall thickness were comparatively greater in older macaques, and did not differ between sexes. In contrast, LVIDd and PWDd demonstrated a sex-difference in both young and older macaques, whereby males had substantially greater dimensions. Sphericity index was similar between older and younger macaques, although females had a significantly greater sphericity index (i.e., a more elongated LV) than males. LV end-diastolic (EDV) and end-systolic volumes (ESV) were comparatively lower in older macaques than their younger counterparts, and were greater in males than in females. Stroke volume did not differ by age, but was greater in males than in females. Sex differences were abolished when LV volumes were scaled to body mass.

*Left ventricular diastolic function;* transmitral early to late diastolic filling velocities (i.e., E/A ratio) was similar between sexes, but was comparatively lower in older macaques compared with their younger counterparts (**Table 3**). Younger females had comparatively larger early filling velocities (E) and late filling velocities (A) than younger males, but sex differences were not present in older macaques. Older males and females had lower E filling velocities than younger macaques; however, only older males had a comparatively greater A filling velocity than their younger counterparts.

**Table 3.** Cardiac function in male and female, young and old free-ranging rhesus macaques.

| Echo Parameter | All | Female |  | Male |  |
| --- | --- | --- | --- | --- | --- |
|  |  | Young<br>(7-12 yrs) | Old<br>(18-26 yrs) | Young<br>(7-12 yrs) | Old<br>(18-26yrs) |
| LV Diastolic Function |  |  |  |  |  |
| Transmitral E/A | 1.7 ± 0.5 | 1.9 ± 0.5 | 1.3 ± 0.4* | 2.0 ± 0.5 | 1.4 ± 0.3* |
| Transmitral E (m/s) | 0.70 ± 0.17 | 0.84 ± 0.12 | 0.64 ± 0.18* | 0.71 ± 0.17† | 0.60 ± 0.10* |
| Transmitral A (m/s) | 0.44 ± 0.13 | 0.47 ± 0.15 | 0.52 ± 0.12 | 0.36 ± 0.08† | 0.45 ± 0.11* |
| Septal e' (mm/s) | 70 ± 16 | 77 ± 15 | 58 ± 12* | 77 ± 12 | 60 ± 17* |
| Septal a' (mm/s) | 48 ± 14 | 49 ± 23 | 50 ± 9 | 42 ± 9 | 57 ± 11* |
| Septal IVRT (ms) | 70 ± 13 | 64 ± 13 | 73 ± 15* | 68 ± 10 | 75 ± 13 |
| Lateral e' (mm/s) | 118 ± 25 | 128 ± 20 | 99 ± 22* | 130 ± 23 | 107 ± 18* |
| Lateral a' (mm/s) | 62 ± 17 | 60 ± 12 | 70 ± 16* | 52 ± 11 | 77 ± 18* |
| Lateral IVRT (ms) | 67 ± 12 | 63 ± 13 | 71 ± 10 | 68 ± 10 | 66 ± 16 |
| LV systolic function |  |  |  |  |  |
| LV Ejection Fraction (%) | 53 ± 5 | 54 ± 4 | 55 ± 4 | 51 ± 5 | 54 ± 3* |
| Septal s' (mm/s) | 51 ± 12 | 58 ± 16 | 48 ± 9* | 50 ± 8 | 47 ± 13 |
| Lateral s' (mm/s) | 75 ± 21 | 82 ± 24 | 74 ± 19 | 69 ± 15 | 76 ± 57 |
| Septal IVCT (ms) | 64 ± 16 | 58 ± 19 | 60 ± 13 | 71 ± 11† | 65 ± 18 |
| Lateral IVCT (ms) | 63 ± 15 | 59 ± 17 | 55 ± 12 | 70 ± 13† | 67 ± 16† |
| LV longitudinal strain (%) | -19.7 ± 2.4 | -20.1 ± 1.9 | -20.4 ± 2.9 | -19.0 ± 2.3 | -19.7 ± 2.7 |
| LV longitudinal strain rate (%/s) | -1.45 ± 0.24 | -1.51 ± 0.12 | -1.47 ± 0.26 | -1.45 ± 0.30 | -1.38 ± 0.20 |
| LA volume and mechanics |  |  |  |  |  |
| LA max volume (ml) | 6 ± 3 | 6 ± 4 | 5 ± 2 | 7 ± 2 | 7 ± 2† |
| LA max volume/body mass (ml/kg) | 0.31 ± 0.07 | 0.30 ± 0.05 | 0.33 ± 0.09 | 0.29 ± 0.06 | 0.33 ± 0.09 |
| LA strain R (%) | 28 ± 6 | 32 ± 7 | 27 ± 6* | 26 ± 5† | 26 ± 4 |
| LA strain CD (%) | -19 ± 6 | -23 ± 5 | -16 ± 5* | -19 ± 5 | -17 ± 5 |
| LA strain CT (%) | -9 ± 4 | -10 ± 3 | -10 ± 5 | -7 ± 3 | -10 ± 5 |

Abbreviations: LV, left ventricle; E/A, ratio of early to late transmitral filling velocity; e', myocardial tissue velocity during early diastole; a' myocardial tissue velocity during late diastole; IVRT, isovolemic relaxation time; IVCT, isovolemic contraction time; LA strain R, left atrial strain during reservoir function; LA strain CD, left atrial strain during conduit function; LA strain CT, left atrial strain during contraction.
\* P < 0.05 vs young, within sex; † P < 0.05 vs female, within age group.

Myocardial tissue velocities during early diastole in the septal (septal e’) and lateral LV walls (lateral e’) were comparatively lower in older than younger macaques, which did not differ by sex. Septal myocardial tissue velocity during late diastole, corresponding with atrial contraction (septal a’) was greater in older males compared with their younger counterparts, but no age difference was observed in females. Tissue velocity of the lateral LV wall during late diastole (lateral a’) was greater in both older males and females in comparison to their younger counterparts. Septal isovolumic relaxation time (IVRT) was significantly greater in older than younger females, but no age effect was observed for males. Additionally, no differences between sexes were observed for septal IVRT or lateral IVRT.

*Left ventricular systolic function;* ejection fraction was similar between males and females, and between older and younger females. However, older males had a greater ejection fraction than younger males. In contrast, systolic myocardial tissue velocity of the septum (septal s’) was significantly lower in older females compared with younger females, while septal s’ was similar between older and younger males. No sex differences were observed in septal s’ or lateral s’, and no age differences were observed for lateral s’. Lateral isovolumic contraction time (IVCT) and septal IVCT did not differ by age group.

However, lateral IVCT was prolonged in males relative to females, for both age groups. Similarly, septal IVCT was greater in younger males compared with their female counterparts, although no sex difference was observed in older macaques. LV longitudinal strain and strain rate did not differ by age or sex.

*Left atrial volumes and mechanics;* LA maximal volume did not differ by age, but was significantly greater in older males compared with older females. However, there were no sex differences when LA maximal volume was scaled to body mass. No sex differences were observed in LA maximal volume (absolute or scaled) in young macaques. A significant interaction effect was found for LA reservoir strain, such that younger females had significantly greater LA reservoir strain in comparison to older females, but older males remained statistically similar to younger males. LA conduit strain did not differ by sex, but was significantly lower in older females compared with younger females, while no difference was observed between young and old males. LA contractile strain did not differ by age or sex.

### The influence of the captive environment on age-related changes in cardiac structure and function

*Population differences in the cardiac phenotype;* on average, free-ranging macaques were older and heavier than their captive counterparts, with a lower heart rate (**Table 4**). Across both captive and free-ranging populations, internal LV dimensions and wall thicknesses were consistently larger in males than in females. However, in comparison to free-ranging macaques, captive individuals had a greater wall thickness (i.e., IVSDd, LVPWd) and hence, a larger relative wall thickness, across both sexes (**Supplementary Table 2**).

**Table 4.** Characteristics of captive and free-ranging rhesus macaque cohorts.

|  | Captive rhesus macaques | Free-ranging rhesus macaques |
| --- | --- | --- |
| Sample size (*n*) | 347 (71% female) | 133 (47% female) |
| Age (years) | 13.2 ± 4.9 | 15.0 ± 3.9* |
| Body mass (kg) | 10.4 ± 3.0 | 8.8 ± 2.0* |
| Heart rate (bpm) | 134 ± 26 | 83 ± 15* |
| **Male** |  |  |
| Age (years) | 12.5 ± 5.5 | 14.8 ± 3.8* |
| Body mass (kg) | 13.5 ± 1.3 | 10.3 ± 1.3* |
| Heart rate (bpm) | 122 ± 22 | 84 ± 14* |
| **Female** |  |  |
| Age | 13.4 ± 4.6 | 15.4 ± 4.0* |
| Body mass | 9.3 ± 2.1<sup>#</sup> | 7.1 ± 0.9* |
| Heart rate (bpm) | 138 ± 27 | 81 ± 15* |

\*Significantly different ( $P < 0.001$ ) from captive rhesus macaques
<sup>#</sup>Significantly different ( $P < 0.001$ ) from male rhesus macaques

Additionally, several diastolic differences were observed between populations. Captive males had a significantly greater E/A ratio than captive females (P = 0.020; **Supplementary Table 2**), but no sex difference was present in free-ranging macaques (P = 0.558). Due to the known influence of heart rate on diastolic function, we also conducted comparisons incorporating both heart rate and age as covariates. Free-ranging macaques had a comparatively greater E/A ratio, when controlling for covariates, than their captive counterparts, in both males (1.76 vs 1.30 in free-ranging vs captive, respectively, P < 0.001; covariates appearing in the model evaluated at age = 13.2 years and heart rate = 98 bpm) and females (1.53 vs 1.25 in free-ranging vs captive, respectively, P < 0.001; covariates appearing in the model evaluated at age = 13.3 and heart rate = 119 bpm). Furthermore, in comparison to captive populations, free-ranging cohorts exhibited significantly greater myocardial tissue velocities during early left ventricular filling (i.e., Avg e’) and less during atrial contraction (Avg a’).

*Left ventricular structure;* relative wall thickness was positively correlated with age in both free-ranging and captive macaques. While the age-related increase in relative wall thickness did not differ between populations within females, captive males had a significantly steeper slope than their free-ranging counterparts (**Figure 1 and Supplementary Table 3 and 4**). This difference between the male populations was not due to LV cavity or wall thickness dimensions alone (i.e., PWD, LVID or IVSd), as slopes for these parameters were similar between free-ranging and captive animals (**Supplementary Table 4**). As such, the population difference in males reflects a combined change in both the LV cavity and wall thickness. Furthermore, the age-related relative wall thickness slope did not differ between sexes within the free-ranging (P = 0.732) and captive populations (P = 0.373).

**Figure 1.**
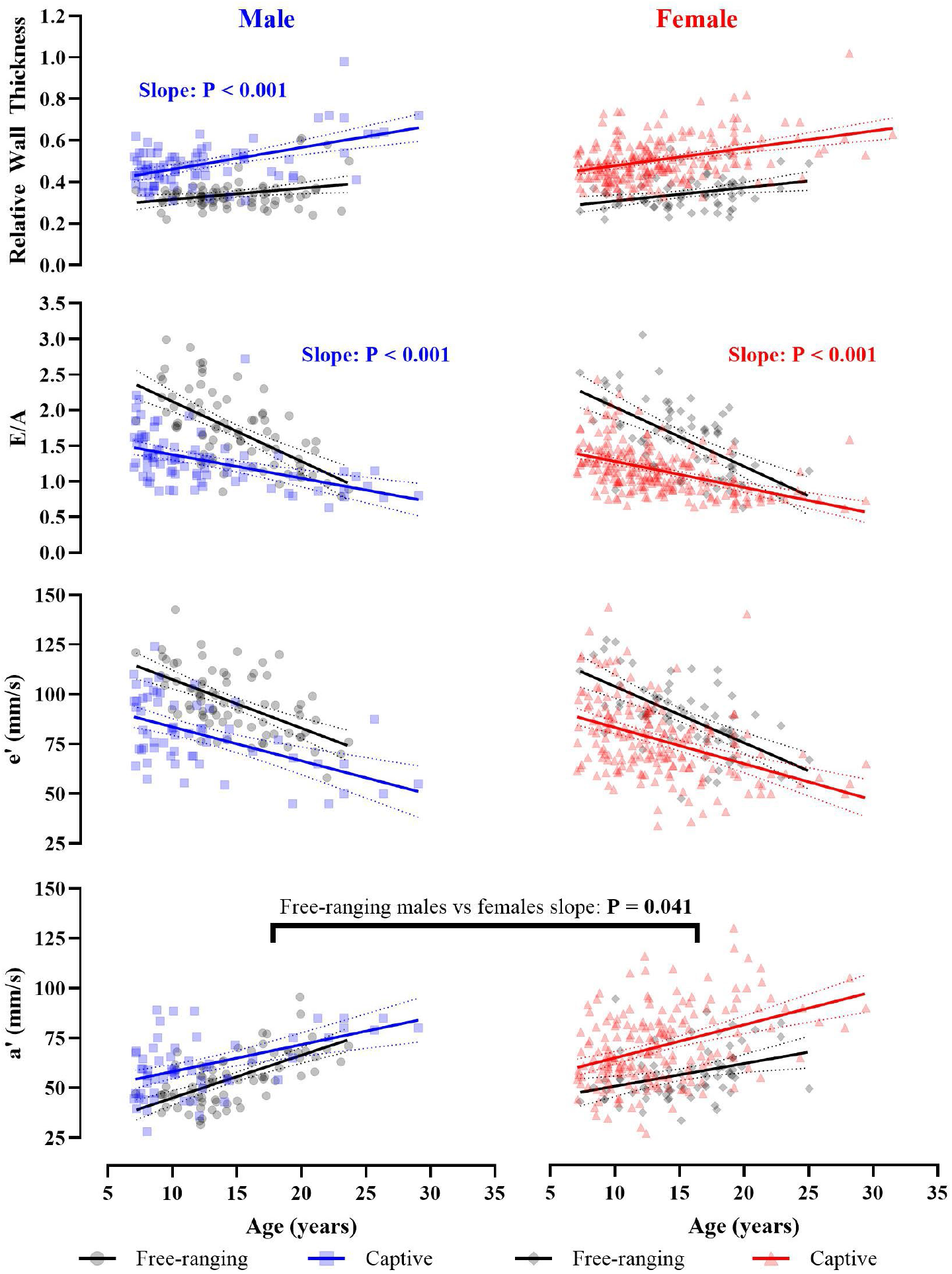
Comparison of age-related slopes of cardiac structure and diastolic function between free-ranging and captive rhesus macaques, showing relative wall thickness, the ratio of the peak velocity of blood flow from left ventricular relaxation in early diastole to late diastole caused by atrial contraction (E/A), peak myocardial tissue velocity during early diastole (e’), and late diastole (a’). Left panels represent comparisons within male macaques (black circles represent free-ranging and blue squares represent captive), while the right panels present comparisons within females (black diamonds represent free-ranging and red triangles represent captive macaques).

*Left ventricular diastolic function;* the relationship between age and E/A ratio was similar between sexes within free-ranging (P = 0.963, Supplementary Table 5) and captive populations (P = 0.818). However, the slope was significantly steeper in free-ranging macaques compared to captive (P < 0.001, **Figure 1**). e′ and a′ were lower in older individuals, with comparable age-related differences across free-ranging and captive populations; however, the slope was significantly greater in free-ranging males compared to free-ranging females (P = 0.041, **Figure 1**).

The interaction effect between age, sex, and environment was P < 0.001 for all parameters. The main effect for environment was P < 0.001 for all parameters. The main effect of sex, and the sex*environment interaction was P > 0.05 for all parameters. P values are provided on the figure where individual slopes are significantly different between sex or environment.

## Discussion

Non-human primates, given their phylogenetic proximity to humans and comparatively short lifespan (7), are an attractive translational model for human cardiovascular aging. However, most research has been derived from captive populations where living conditions are markedly different from natural habitats (9,16,19–21). Captive settings offer limited space, absent or highly contrived social networks, and environmental conditions, such as temperature and light exposure, that do not reflect natural habitats (22) - all of which may impact cardiovascular outcomes. In this study, we provide the first comprehensive characterization of cardiac health across age in free-ranging rhesus macaques, in the absence of artificial confounders. As predicted, we found that greater chronological age is associated with thickening of left ventricular walls relative to cavity diameter, and lower diastolic function in free-ranging macaques. Furthermore, at a comparable stage of life, captive animals demonstrated a more aged cardiac phenotype relative to their free-ranging counterparts. This highlights the critical role of environment and lifestyle in shaping cardiac health across age.

As expected, we observed hallmark features of cardiac aging in older free-ranging rhesus macaques. These included smaller LV volumes, greater relative wall thickness, and lower systolic and diastolic function in older animals. Furthermore, males exhibited greater wall thicknesses, diameters, and volumes in comparison to females. Yet, when normalized to body mass, differences were no longer evident, indicating that these apparent sex differences primarily reflect body size rather than true sexual dimorphism. Interestingly, females have a more elongated left ventricle, as indicated by a greater sphericity index. While the functional significance of this sex-specific pattern is not fully understood, an elongated shape has been associated with enhanced LV twist mechanics (23,24), possibly due to more optimal myofiber architecture facilitating more efficient function. In addition, while diastolic function (as measured via transmitral E/A ratio) was similar between sexes, females showed higher absolute diastolic filling velocities in comparison to males. It is possible that the sex differences in geometry and diastolic function reflect adaptations to the varying hemodynamic demands associated with sex-specific life histories. To examine this, an understanding of the cardiac phenotype during critical phases of life in males and females is required.

While age-related differences have previously been observed in captive macaques (9), and are now evident in a free-ranging population, differences between these environments may result in distinct aging patterns.

In line with our predictions, we show that hallmark features of cardiac aging are significantly different between captive and free-ranging animals. Specifically, irrespective of age or sex, captive animals present with a greater relative wall thickness in comparison to free-ranging macaques. In humans, a relative wall thickness greater than 0.42 would be indicative of concentric remodeling (17), often associated with pathology, such as hypertensive heart disease (25,26). While free-ranging animals were largely below 0.42 (∼ 0.34), captive animals far exceeded the human threshold (∼ 0.50). However, whether this difference in structure reflects a genetic or acquired phenotype is not clear; but, given the known influence of physical inactivity, poor diet, and social stress on human cardiovascular health, it is likely that these factors contribute to the thicker ventricular walls seen in this study and other captive macaque populations (9,16,19). Indeed, Florio et al., (9) observed no difference in relative wall thickness between young and old captive rhesus macaques, suggesting that lifestyle and environmental influences may begin early in life, potentially predisposing individuals to adverse cardiac outcomes later in life.

In humans, age-related changes in ventricular structure are often accompanied by a decline in diastolic function (27). Ventricular wall thickening and associated changes in myocardial ultrastructure result in a stiffening of the left ventricle, meaning that compliance and early diastolic recoil are impaired (28,29). In line with this, older individuals exhibited lower diastolic function (E/A ratio and e’) across both free-ranging and captive animals. However, our data indicate that young free-ranging animals have greater diastolic function (i.e., a greater E/A ratio, higher e’ and lower a’) than their captive counterparts; yet intriguingly, by ∼25 years of age, the age-related differences converge (E/A ratio in both males and females, and a’ in males only; **Figure 1**). It is possible that young captive animals have a comparatively lower diastolic function that results from adverse maternal programming related to artificial environmental conditions (e.g., epigenetic modification or poor intrauterine environment) (30,31), early life adversity (e.g. maternal separation or abnormal social dynamics) (32–34), and/or the direct impact of captivity (e.g. limited space for physical activity, poor nutrition, chronic psychological stress) (35). Despite the potentially negative impact of the captive environment, captive animals frequently live longer than free-ranging (7). It is possible that 25 years of age reflects a critical threshold in macaques, whereby diastolic function is unable to meet the physiological demands of the naturalistic environment. Human intervention (i.e., provisioning of resources and care), as occurs in captivity, may artificially prolong the lifespan, which could explain discrepancies in the age-related slopes between captive and free-ranging populations (**Figure 1**). It is also possible that these discrepancies may reflect differences in allostatic load (i.e., cumulative burden of stress and life events across physiological systems) (36) or survivorship bias in our cross-sectional cohorts. Future research should integrate longitudinal, multi-system approaches to disentangle genetic, social, and environmental contributions to cardiovascular aging.

### Limitations

Although free-ranging macaques inhabit a naturalistic environment, they are not exposed to all elements of a truly wild habitat which may influence the generalizability of our findings. This, however, creates a model that is more similar to the modern niche of many human populations, characterized by access to abundant nutritional resources and a lower risk of predation than is present in the wild. Additionally, while the environmental contrast between free-ranging and captive macaques provides a valuable model for examining lifestyle effects, the cross-sectional design limits our ability to establish causality or track individual aging trajectories over time. Lastly, while all animals were handled in accordance with best-practice veterinary protocols, different anesthetic protocols were used in free-ranging and captive populations. Although all protocols were designed to minimize physiological disturbance, we cannot fully exclude the possibility that anesthetic differences may have influenced measures of cardiac function. However, the influence of anesthesia on cardiac structure is likely to be minimal.

## Conclusion

In conclusion, our findings provide the first comprehensive characterization of age-related cardiac remodeling in free-ranging rhesus macaques, revealing structural and functional changes that parallel hallmark features of human cardiovascular aging. Differences between captive and free-ranging macaques were apparent even in young adults, suggesting that hereditary factors and the developmental environment are likely important determinants of the adult cardiac phenotype. Thus, our data highlight the importance of the ecological context when interpreting data from animal models. It is possible that captive models present a distorted view of normal physiology, whereby the living environment differs markedly from the ancestral environment. While such models remain valuable for exploring the physiological consequences of aging, it is crucial that they are framed and interpreted within this ecological and evolutionary context.

## Acknowledgements

The authors thank all staff at the Caribbean Primate Research Facility and the California National Primate Research Centre for their support in data collection.

