## Supplementary Information for "The influence of captivity on cardiac structure and function across age, in rhesus macaques"

**Supplementary Table 1**. ANOVA P values for main effects and pairwise comparisons of cardiac structure and function in male and female young and old free-ranging rhesus macaques (extension of Table 2 in manuscript)

| Parameter | Interaction | Main effect | | Age Pairwise comparison | | Sex Pairwise comparison | |
|---|---|---|---|---|---|---|---|
| | Age*sex<br>P value | Age<br>P value | Sex<br>P value | Young vs old male<br>P value | Young vs old female<br>P value | Young Male vs female<br>P value | Old male vs female<br>P value |
| LV mass (g) | 0.363 | 0.291 | **< 0.001** | | | **< 0.001** | **< 0.001** |
| LV mass/body mass (g/kg$^{1.0}$) | 0.247 | **< 0.001** | 0.808 | **0.001** | 0.112 | | |
| Aortic annulus diameter (mm) | 0.385 | **0.004** | **< 0.001** | **0.007** | 0.150 | **< 0.001** | **< 0.001** |
| Aortic annulus diameter/body mass (mm/kg$^{0.25}$) | 0.275 | **< 0.001** | **< 0.001** | **< 0.001** | **0.005** | **0.022** | **< 0.001** |
| IVSDd (mm) | 0.731 | **< 0.001** | 0.123 | **0.002** | **0.033** | | |
| IVSDd/body mass (mm/kg$^{0.25}$) | 0.735 | **< 0.001** | 0.790 | **0.002** | **0.011** | | |
| LVIDd (mm) | 0.129 | 0.062 | **< 0.001** | | | **< 0.001** | **< 0.001** |
| LVIDd/body mass (mm/kg$^{0.25}$) | 0.072 | 0.585 | **0.016** | | | 0.626 | **0.006** |
| PWDd (mm) | 0.582 | 0.486 | **0.002** | | | **0.004** | **0.004** |
| PWDd/body mass (mm/kg$^{0.25}$) | 0.716 | 0.113 | 0.452 | | | | |
| RWT | 0.285 | **0.001** | 0.304 | 0.116 | **0.003** | | |
| Sphericity index | 0.839 | 0.241 | **0.006** | | | **0.050** | **0.046** |
| **LV Volumes** | | | | | | | |
| LV EDV biplane (ml) | 0.801 | **< 0.001** | **< 0.001** | **0.004** | **0.016** | **< 0.001** | **< 0.001** |
| LV EDV/body mass (ml/kg$^{1.0}$) | 0.724 | 0.102 | 0.087 | | | | |
| LV ESV biplane (ml) | 0.279 | **< 0.001** | **< 0.001** | **< 0.001** | 0.030 | **< 0.001** | **0.001** |
| LV ESV/body mass (ml/kg$^{1.0}$) | 0.849 | **0.022** | 0.217 | 0.072 | 0.142 | | |
| LV SV biplane (ml) | 0.670 | 0.066 | **< 0.001** | | | **0.001** | **0.001** |
| LV SV/body mass (ml/kg$^{1.0}$) | 0.663 | 0.677 | 0.091 | | | | |
| **LV Diastolic Function** | | | | | | | |
| E/A | 0.942 | **< 0.001** | 0.427 | **< 0.001** | **< 0.001** | | |
| Transmitral E (m/s) | 0.207 | **< 0.001** | **0.013** | **0.020** | **< 0.001** | **0.004** | 0.401 |
| Transmitral A (m/s) | 0.354 | **0.006** | **0.001** | **0.008** | 0.182 | **0.002** | 0.120 |
| Septal e’ (mm/s) | 0.799 | **< 0.001** | 0.716 | **< 0.001** | **< 0.001** | | |
| Septal a’ (mm/s) | **0.025** | **0.010** | 0.957 | **< 0.001** | 0.803 | 0.075 | 0.145 |
| Septal IVRT (ms) | 0.692 | **0.006** | 0.330 | 0.090 | **0.028** | | |
| Lateral e’ (mm/s) | 0.540 | **< 0.001** | 0.314 | **< 0.001** | **< 0.001** | | |
| Lateral a’ (mm/s) | **0.021** | **< 0.001** | 0.893 | **< 0.001** | **0.031** | 0.059 | 0.147 |
| Lateral IVRT (ms) | 0.105 | 0.300 | 0.967 | | | | |
| **LV systolic function** | | | | | | | |
| LV EF Biplane (%) | 0.273 | **0.032** | 0.075 | **0.019** | 0.459 | | |
| Septal s’ (mm/s) | 0.113 | **0.019** | 0.068 | 0.569 | **0.006** | | |
| Lateral s’ (mm/s) | 0.098 | 0.944 | 0.255 | | | | |
| Septal IVCT (ms) | 0.250 | 0.624 | **0.010** | | | **0.004** | 0.331 |
| Lateral IVCT (ms) | 0.914 | 0.270 | **< 0.001** | | | **0.009** | **0.018** |
| LV longitudinal strain (%) | 0.760 | 0.404 | 0.167 | | | | |
| LV longitudinal strain rate (%/s) | 0.842 | 0.410 | 0.231 | | | | |
| **LA volume and mechanics** | | | | | | | |
| LA max volume (ml) | 0.493 | 0.374 | **0.016** | | | 0.194 | 0.035 |

| | | | | | | | |
|---|---|---|---|---|---|---|---|
| LA max volume/body mass (ml/kg) | 0.885 | 0.057 | 0.986 | | | | |
| LA strain R (%) | **0.029** | 0.069 | 0.026 | 0.778 | **0.006** | **0.002** | 0.973 |
| LA strain CD (%) | 0.100 | **0.002** | 0.305 | 0.272 | **0.001** | | |
| LA strain CT (%) | 0.265 | 0.065 | 0.097 | | | | |

**Supplementary Table 2.** ANCOVA comparing cardiac structure and diastolic function between male and female rhesus macaques living in captive and free-ranging environments, using age as a covariate.

| | **Captive** | | | **Free-ranging** | | | | | **Main effect:** |
|---|---|---|---|---|---|---|---|---|---|
| | **Male** | **Female** | **Sex difference in captivity (P value)** | **Male** | **Female** | **Sex difference in free-ranging (P value)** | **Male Captive vs free-ranging (P value)** | **Female Captive vs free-ranging (P value)** | **i) Age*Env*Sex interaction ii) Env*Sex interaction iii) Environment iv) Sex (P value)** |
| **RWT** | | | | | | | | | i) < **0.001** |
| Mean ± SD | 0.49 ± 0.11 | 0.51 ± 0.10 | | 0.33 ± 0.06 | 0.35 ± 0.07 | | **< 0.001** | **< 0.001** | ii) 0.952 |
| adj. Mean ± SE | 0.49 ± 0.01 | 0.51 ± 0.01 | | 0.33 ± 0.01 | 0.34 ± 0.01 | | | | iii) **< 0.001** |
| | | | | | | | | | iv) 0.155 |
| **IVSd (mm)** | | | | | | | | | i) < **0.001** |
| Mean ± SD | 5.8 ± 0.9 | 5.1 ± 1.0 | **< 0.001** | 3.9 ± 0.8 | 3.7 ± 0.8 | 0.264 | **< 0.001** | **< 0.001** | **ii) 0.002** |
| adj. Mean ± SE | 5.9 ± 0.1 | 5.1 ± 0.1 | | 3.8 ± 0.1 | 3.7 ± 0.1 | | | | iii) **< 0.001** |
| | | | | | | | | | iv) <**0.001** |
| **LVIDd (mm)** | | | | | | | | | i) **< 0.001** |
| Mean ± SD | 25.9 ± 4.2 | 21.9 ± 2.9 | **< 0.001** | 24.2 ± 2.0 | 22.4 ± 2.4 | **< 0.001** | **0.004** | 0.106 | ii) < **0.001** |
| adj. Mean ± SE | 25.7 ± 0.3 | 21.9 ± 0.2 | | 24.3 ± 0.4 | 22.6 ± 0.4 | | | | iii) 0.256 |
| | | | | | | | | | iv) **< 0.001** |
| **LVPWd (mm)** | | | | | | | | | i) **< 0.001** |
| Mean ± SD | 6.4 ± 0.7 | 5.8 ± 0.9 | **< 0.001** | 4.1 ± 0.6 | 4.0 ± 0.7 | **0.002** | **< 0.001** | **< 0.001** | ii) **0.009** |
| adj. Mean ± SE | 6.4 ± 0.08 | 5.8 ± 0.05 | | 4.1 ± 0.1 | 3.9 ± 0.1 | | | | iii) **< 0.001** |
| | | | | | | | | | iv) **< 0.001** |
| **E/A** | | | | | | | | | i) **< 0.001** |
| Mean ± SD | 1.30 ± 0.37 | 1.15 ± 0.33 | **0.020** | 1.71 ± 0.48 | 1.65 ± 0.52 | 0.558 | **< 0.001** | **< 0.001** | ii) 0.355 |
| adj. Mean ± SE | 1.23 ± 0.04 | 1.13 ± 0.02 | | 1.75 ± 0.04 | 1.71 ± 0.04 | | | | iii) **< 0.001** |
| | | | | | | | | | iv) 0.063 |
| **E (mm/s)** | | | | | | | | | i) **< 0.001** |
| Mean ± SD | 0.74 ± 0.19 | 0.72 ± 0.16 | | 0.68 ± 0.15 | 0.72 ± 0.16 | | | | ii) 0.183 |
| adj. Mean ± SE | 0.72 ± 0.02 | 0.71 ± 0.01 | | 0.69 ± 0.02 | 0.73 ± 0.02 | | | | iii) 0.689 |
| | | | | | | | | | iv) 0.414 |
| **A (mm/s)** | | | | | | | | | i) **< 0.001** |
| Mean ± SD | 0.58 ± 0.14 | 0.64 ± 0.17 | **0.002** | 0.42 ± 0.11 | 0.46 ± 0.12 | 0.208 | **< 0.001** | **< 0.001** | ii) 0.385 |
| adj. Mean ± SE | 0.59 ± 0.02 | 0.65 ± 0.01 | | 0.42 ± 0.02 | 0.48 ± 0.02 | | | | iii) **< 0.001** |
| | | | | | | | | | iv) **0.004** |
| **Avg e' (mm/s)** | | | | | | | | | i) **< 0.001** |
| Mean ± SD | 79 ± 21 | 79 ± 19 | | 93 ± 18 | 91 ± 18 | | **< 0.001** | **< 0.001** | ii) 0.579 |
| adj. Mean ± SE | 77 ± 2 | 79 ± 1 | | 94 ± 2 | 93 ± 2 | | | | iii) **< 0.001** |
| | | | | | | | | | iv) 0.966 |
| **Avg a' (mm/s)** | | | | | | | | | i) **< 0.001** |
| Mean ± SD | 65 ± 14 | 70 ± 17 | | 56 ± 15 | 55 ± 12 | | **< 0.001** | **< 0.001** | ii) 0.063 |
| adj. Mean ± SE | 66 ± 2 | 70 ± 1 | | 56 ± 2 | 54 ± 2 | | | | iii) **< 0.001** |
| | | | | | | | | | iv) 0.376 |

RWT, relative wall thickness; IVSd, interventricular septal diameter in diastole; LVIDd, left ventricular internal diameter in diastole; LVPWd, left ventricular posterior wall thickness in diastole; E/A, ratio of early to later left ventricular filling velocity; E, early left ventricular filling velocity; A, late left ventricular filling velocity; Avg e', an average of myocardial tissue velocity during early left ventricular filling taken at the septal and lateral wall; Avg a', an average of myocardial tissue velocity during late left ventricular filling taken at the septal and lateral wall. Age covariate appearing in the model at age = 13.7

**Supplementary Table 3**. Slopes and intercepts of the relationship between cardiac variables and age in captive and free-ranging male and female rhesus macaques.

| | Captive | | Free-ranging | |
|---|---|---|---|---|
| **Parameter** | **Male** | **Female** | **Male** | **Female** |
| **RWT** | | | | |
| Slope | 0.011 | 0.008 | 0.005 | 0.006 |
| Intercept | 0.358 | 0.395 | 0.225 | 0.246 |
| P value | **< 0.001** | **< 0.001** | **0.012** | **0.004** |
| **IVSd** | | | | |
| Slope | 0.034 | 0.052 | 0.073 | 0.047 |
| Intercept | 5.47 | 4.48 | 2.91 | 2.88 |
| P value | 0.071 | **< 0.001** | **0.005** | **0.035** |
| **LVIDd** | | | | |
| Slope | -0.244 | -0.148 | -0.118 | -0.139 |
| Intercept | 29.0 | 24.1 | 26.1 | 24.2 |
| P value | **0.004** | **< 0.001** | 0.055 | **0.009** |
| **LVPWd** | | | | |
| Slope | 0.058 | 0.037 | 0.003 | 0.037 |
| Intercept | 5.67 | 5.36 | 4.26 | 3.33 |
| P value | **< 0.001** | **< 0.001** | 0.896 | 0.119 |
| **E/A** | | | | |
| Slope | -0.033 | -0.037 | -0.084 | -0.083 |
| Intercept | 1.71 | 1.64 | 2.96 | 2.87 |
| P value | **< 0.001** | **< 0.001** | **< 0.001** | **< 0.001** |
| **E** | | | | |
| Slope | -0.013 | -0.010 | -0.015 | -0.021 |
| Intercept | 0.897 | 0.850 | 0.904 | 1.06 |
| P value | **< 0.001** | **< 0.001** | **< 0.001** | **< 0.001** |
| **A** | | | | |
| Slope | 0.005 | 0.009 | 0.013 | 0.012 |
| Intercept | 0.513 | 0.522 | 0.226 | 0.315 |
| P value | **0.045** | **< 0.001** | **< 0.001** | **0.002** |
| **Avg e’** | | | | |
| Slope | -2.44 | -1.82 | -2.44 | -2.82 |
| Intercept | 29.0 | 101 | 132 | 132 |
| P value | **< 0.001** | **< 0.001** | **< 0.001** | **< 0.001** |
| **Avg a’** | | | | |
| Slope | 2.14 | 1.66 | 2.14 | 1.14 |
| Intercept | 23.5 | 48.4 | 23.5 | 39.3 |
| P value | **< 0.001** | **< 0.001** | **< 0.001** | **0.004** |
| **Body mass** | | | | |
| Slope | 0.079 | 0.056 | -0.018 | -0.079 |
| Intercept | 12.4 | 8.59 | 10.5 | 8.37 |
| P value | 0.194 | 0.055 | 0.446 | **0.004** |

**Supplementary Table 4.** Comparison of slopes and intercepts of the relationship between cardiac variables and age between male and female captive and free-ranging rhesus macaques.

| | Captive<br><br>Male vs female | Free ranging<br><br>Male vs female | Males<br><br>Free-ranging vs captive | Females<br><br>Free-ranging vs captive |
|---|---|---|---|---|
| **RWT** | | | | |
| **Slopes** | P = 0.373 | P = 0.732 | **P < 0.001** | P = 0.521 |
| **Intercepts** | P = 0.349 | P = 0.862 | | **P < 0.001** |
| **IVSd** | | | | |
| **Slopes** | P = 0.446 | P = 0.455 | P = 0.232 | P = 0.960 |
| **Intercepts** | **P < 0.001** | **P = 0.002** | P < **0.001** | P < **0.001** |
| **LVIDd** | | | | |
| **Slopes** | P = 0.188 | P = 0.793 | P = 0.287 | P = 0.892 |
| **Intercepts** | **P < 0.001** | P = < **0.001** | P = **0.037** | P = 0.201 |
| **LVPWd** | | | | |
| **Slopes** | P = 0.294 | P = 0.245 | P = 0.306 | P = 0.837 |
| **Intercepts** | **P < 0.001** | **P < 0.001** | **P < 0.001** | **P < 0.001** |
| **E/A** | | | | |
| **Slopes** | P = 0.818 | P = 0.963 | P < **0.001** | P < **0.001** |
| **Intercepts** | P = **0.001** | P = 0.236 | | |
| **E** | | | | |
| **Slopes** | P = 0.479 | P = 0.425 | P = 0.667 | P = 0.056 |
| **Intercepts** | P = 0.641 | P = **0.002** | P = 0.279 | P = **0.028** |
| **A** | | | | |
| **Slopes** | P = 0.317 | P = 0.792 | P = 0.088 | P = 0.672 |
| **Intercepts** | P = **0.004** | P = **0.001** | **P < 0.001** | **P < 0.001** |
| **e'** | | | | |
| **Slopes** | P = 0.817 | P = 0.526 | P = 0.175 | P = 0.094 |
| **Intercepts** | P = 0.778 | P = **0.014** | **P < 0.001** | **P < 0.001** |
| **a'** | | | | |
| **Slopes** | P = 0.519 | P = **0.041** | P = 0.074 | P = 0.370 |
| **Intercepts** | **P = 0.002** | | **P < 0.001** | **P < 0.001** |
